# Establishment of primary transgenic human airway epithelial cell cultures to study respiratory virus – host interactions

**DOI:** 10.1101/694380

**Authors:** Hulda R. Jonsdottir, Sabrina Marti, Dirk Geerts, Regulo Rodriguez, Volker Thiel, Ronald Dijkman

**Author notes:** Correspondence: Ronald Dijkman, Institute of Virology and Immunology, Department of infectious Diseases and Pathobiology, Vetsuisse Faculty, University of Bern, Länggassstrasse 122, 3012 Bern, Switzerland. Institute of Microbiology, Lausanne University Hospital and University of Lausanne, Lausanne, Switzerland. Contributed equally (Volker Thiel, Ronald Dijkman).

## Abstract

Primary human airway epithelial cell (hAEC) cultures represent a universal platform to propagate respiratory viruses and characterize their host interactions in authentic target cells. To further elucidate specific interactions between human respiratory viruses and important host factors in airway epithelium, it is important to make hAEC cultures amenable to genetic modification. However, the short and finite lifespan of primary cells in cell culture creates a bottleneck for the genetic modification of these cultures. In the current study, we show that the incorporation of the Rho-associated protein kinase (ROCK) inhibitor (Y-27632) during cell propagation extends the life span of primary human cells in vitro and thereby facilitates the incorporation of lentivirus-based expression systems. Using fluorescent reporters for FACS-based sorting, we generated homogenously fluorescent hAEC cultures that differentiate normally after lentiviral transduction. As proof-of-principle, we demonstrate that host gene expression can be modulated post-differentiation via inducible short hairpin (sh)RNA-mediated knockdown. Importantly, functional characterization of these transgenic hAEC cultures with exogenous poly(I:C), as a proxy for virus infection, demonstrates that such modifications do not influence the host innate immune response. Moreover, the propagation kinetics of both human coronavirus 229E (HCoV-229E) and human respiratory syncytial virus (RSV) were not affected. Combined, these results validate our newly established protocol for the genetic modification of hAEC cultures thereby unlocking a unique potential for detailed molecular characterization of virus – host interactions in human respiratory epithelium.

## 1. Introduction

The human lungs are a large organ and span a relatively long anatomical distance. As a result, pulmonary histology differs substantially depending on anatomical location and specific tissue function. The upper airways are ciliated, pseudostratified and contain multiple cell types with varying roles in the differentiated tissue [1]. Goblet cells produce protective mucus, ciliated cells are responsible for cleaning out both mucus and debris [2], while basal cells serve as resident progenitor cells and replenish other cell types [3].

Human airway epithelial cell (hAEC) cultures are organotypic air-liquid interface (ALI) cell cultures that morphologically and functionally resemble the human airway in vivo [4]. Currently, both the upper and lower airways can be recapitulated in vitro by using this cell culture system. To establish ALI cell cultures, the cells are cultured on porous inserts where only the basolateral side is in contact with growth medium while the apical side is exposed to air, resembling the orientation of in vivo airway epithelium. To represent different areas of the pulmonary epithelium, different cell types can be cultured in the system. Tracheobronchial hAEC cultures contain, after differentiation, different cell types, including basal, ciliated and goblet cells. Moreover, these cultures are pseudostratified and generate protective mucus much like in vivo tracheobronchial epithelium [5,6]. As a result, such cultures are ideal for virus - host interaction studies with human respiratory viruses since they represent the primary entry point of these pathogens [7–13]. Traditionally, virus – host interactions are studied in animal models and human respiratory viruses are usually investigated in ferrets and transgenic mice [14,15]. However, in order to infect these animals, viruses often have to be adapted to the animal by serial passage and this may cause both genotypic and phenotypic differences between the original human virus and the adapted one. Furthermore, it is often difficult to translate results obtained in animal models directly to human disease. Therefore, it is important to study human viruses in authentic human target cells. We, and others, have previously demonstrated that hAEC cultures serve as a universal platform for the study and propagation of human respiratory viruses [7–13].

However, in order to fully utilize the potential of this culture system it must be made amenable to genetic modification. Transgenesis would enable the study of viral and/or host factors important for respiratory virus infections and allow for the elucidation of specific mechanism involved in virus-host interactions by targeted gene knockdown or overexpression. However, using primary cells for genetic modification is challenging since they have a finite life span in cell culture. Primary human bronchial cells can only be limitedly passaged after isolation if differentiation capabilities are to be maintained [16]. The incorporation of the Rho-associated protein kinase (ROCK) inhibitor Y-27632 has been shown to increase the number of passages primary cells can undergo in vitro without gross influence on cell differentiation capacity [17,18]. Theoretically, this would enable the generation of genetically modified well-differentiated primary human airway epithelium in vitro.

Due to the extended culture time required for hAEC culture establishment and differentiation, and the fact that well-differentiated cultures can be maintained for months [19], stable integration of any genes to be expressed post-differentiation is required. Therefore, we established our protocol using lentiviral vectors, where the proviral genomic material is integrated directly into the cellular genome. Furthermore, such a system allows for the integration of both transgene and shRNA scaffolds for over expression and/or knockdown of both host and viral factors [20–22]. Although protocols for the generation of transgenic airway epithelium for various purposes have been described [23–25], thus far there have been no reports describing whether lentivirus modification of well-differentiated hAECs alters the host innate immune response and/or susceptibility to viral infection.

In order to establish homogeneously transgenic hAEC cultures suitable for characterization of virus – host interactions we adapted our current cell culture protocol to generate genetically modified well-differentiated hAECs with uniform transgene distribution. As proof-of-principle, we assessed whether the host gene expression can be modulated post-differentiation via inducible short hairpin (sh)RNA-mediated knockdown and determined whether this affected the innate immune response or the susceptibility to viral infection by two common cold viruses.

## 2. Materials and Methods

### 2.1 Cell culture

The human cell lines 293LTV (LTV-100; Cellbiolabs, San Diego, California, USA) and Huh-7 (gift from V. Lohmann) were maintained in Dulbecco’s Modified Eagle Medium-GlutaMAX supplemented with 1 mM sodium pyruvate, 10% heat-inactivated fetal bovine serum, 100 µg/ml Streptomycin, 100 IU/ml Penicillin and 0.1 mM MEM Non-Essential Amino Acids (Gibco; Thermo Fisher Scientific). All cell lines were culture at 37°C in a humidified incubator with 5% CO_2_.

### 2.2 Human airway epithelial cell culture

Primary human tracheobronchial cells were isolated from patients (>18 years old) undergoing bronchoscopy or pulmonary resection at the Cantonal Hospital in St. Gallen, Switzerland, in accordance with our ethical approval (EKSG 11/044, EKSG 11/103 and KEK-BE 302/2015). Isolation of cells was performed with protease and DNase digestion and primary human tracheobronchial cells were cultured as previously described [19], with the following modifications. Monolayer cultures of primary tracheobronchial epithelial cells were cultured on collagen-coated flasks in complete Bronchial Epithelial Growth Medium (BEGM) supplemented with 100 μg/ml Streptomycin, 100 IU/ml Penicillin, and with or without 10 μM Rho-associated protein kinase inhibitor (Y-27632, Abcam, Cambridge, United Kingdom). For establishment of well-differentiated airway epithelial cell cultures, primary human tracheobronchial epithelial cells were seeded in BEGM with or without 10 μM Y-27635 onto Collagen Type IV coated 24-well Transwell HTS plates (Corning) at the density of 250.000 cells per cm^2^. Once cells reached confluency, the apical and basolateral medium was changed to air-liquid interphase (ALI) medium, and one day later the apical side medium was aspirated, establishing ALI. The epithelial layer was allowed to differentiate for at least four weeks prior to any analyses. Basolateral medium was changed every other day during the differentiation period. Additionally, the cell layer was washed with 200 µL Hank’s Balanced Salt Solution (HBSS, Gibco; Thermo Fisher Scientific) once a week.

### 2.3 Lentiviral vectors

Third-generation lentiviral packaging plasmids (pMDL (gagPol), pRev, pVSV-g) were generously provided by Prof. Dr. B. Berkhout and Dr. J. Eekels along with the eGFP-modified pLKO backbone containing a non-mammalian short-hairpin (sh)RNA control (pLKO_GFP; Mission™ SHC202) that should not target any known mammalian gene, but will engage with RISC [26]. The Isopropyl β-D-1-thiogalactopyranoside (IPTG)-inducible lentiviral backbone pLKO-puro-IPTG_3xLacO harboring the non-mammalian shRNA control was purchased (Mission™SHC332; Sigma Aldrich, Buchs, Switzerland). The puromycin resistance gene was replaced via restriction digestion cloning using a PCR-amplified fluorescent mCherry marker containing overhangs with compatible restriction sites to generate the pLKO-mCherry_IPTG_3xLacO lentiviral backbone.

### 2.4 shRNA ligation

The shRNA sequences TRCN0000231750 and TRCN0000231753 for knockdown of eGFP were extracted from the RNAi consortium website (https://www.broadinstitute.org/rnai/public/) and ordered as separate sense and antisense strands (Microsynth AG, Balgach, Switzerland) incorporating specific restriction sites. shRNA oligos were annealed in CutSmart buffer (New England Biolabs) by incubation of the sense and antisense strands at 94°C for 10 minutes and slow cooling to room temperature (RT). The generated double stranded shRNAs were inserted into the pLKO-mCherry_IPTG_3xLacO lentiviral backbone via restriction digestion cloning.

### 2.5 Lentiviral particle production

Low passage 293LTV cells were seeded in 2.2 mL of culture medium in a T25 cell culture flask (TPP, Trasadingen, Switzerland) at a density of 880.000 cells/cm2 16 - 18 hours prior to transfection. For transfection, a total of 2.4 µg transfer vector, 1.6 µg pMDL, 0.6 µg pRev and 0.8 µg pVSV-g was transfected into the cells using Lipofectamine 2000® (Fisher Scientific AG, Reinach, Switzerland) according to the manufacturer protocol. Twenty-four hours post-transfection, the medium was replaced with BEGM and incubated for an additional 24 – 48 hours before lentivirus-containing supernatant was collected on ice and spun down at 400 *x rcf* for 5 minutes at 4°C. Lentiviral titer was estimated using the GoStix rapid lentiviral titer detection kit (Takara Bio Europe SAS, Saint-Germain-en-Laye, France). Lentiviruses were either used directly for transduction of primary tracheobronchial cells or stored at −80°C.

### 2.6 Lentiviral transduction

Undifferentiated primary human tracheobronchial cells were transduced in suspension with 500 μL lentiviral supernatant for 4 hours at 37°C in batches of 100.000 cells in 1 mL total BEGM, supplemented with 10 μM Y-27635, with gentle shaking every hour. Subsequently, cells were seeded into T25 flasks (TPP) for monolayer culture in 4 mL total medium with lentiviral supernatant for 24 hours prior to washing with HBSS and cell maintenance as described above. Control cells were incubated accordingly to account for any experimental effects. Once confluent, cells were expanded into T75 flasks (TPP).

### 2.7 Flow Cytometry

Cells were trypsinized with 0.05% Trypsin/EDTA (Gibco), resuspended and fixed with 1 mL 4% buffered formalin (FORMAFIX, Formafix Switzerland AG, Hittnau, Switzerland) at RT for 15 minutes and washed with PBS (400 *x rcf*, 5 min, 4°C). Cells were stained with antibodies against tubulin (3624S, Alexa Fluor-488; Cell Signaling, Bioconcept AG, Allschwil, Switzerland), Nerve growth factor receptor (NGFR, 562122, PE-Cy7; BD Bioscience) and Mucin 1 (355604, PE; Biolegend, London, United Kingdom) in 100 μL Cell Wash buffer (CWB, BD Bioscience, Allschwil, Switzerland) in batches of 200.000 cells on ice for 20 minutes and washed twice in 1 mL CWB (400 *x rcf*, 5 min, 4°C) Cells were then resuspended in 100μL of CWB and analyzed with FACS Canto (BD Bioscience). For quantification of GFP expression, cells were analyzed by flow cytometry directly. Prior to analysis cells were fixed as described above and subsequently washed with HBSS. Cells were then resuspended in HBSS and analyzed with FACS Canto using non-transduced cells as negative control.

### 2.8 FACS sorting

After lentiviral transduction and monolayer expansion, transduced cells were sorted for single positive mCherry signal or double positive mCherry/eGFP signal at 4°C using FACS Aria III and the corresponding FACS Diva software (BD Bioscience). Cells were sorted from HBSS supplemented with 10 µM Y-27632, and 0.1% Pluronic (Sigma Aldrich) into BEGM supplemented with 10 µM Y-27632 in FACS flow (BD Bioscience) and washed with HBSS (400 *x rcf*, 5 min, 4°C) prior to further culturing. Cells were resuspended in complete BEGM supplemented with 10 µM Y-27632, amphotericin B and gentamicin (Sigma Aldrich). Medium was changed to complete BEGM supplemented with 10 µM Y-27632 the next day and every other day thereafter until 90% confluency was reached. Cells were then expanded to larger culture flasks. Cells were sorted at the FACS core facility, Institute of Pathology, University of Bern, Bern, Switzerland.

### 2.9 Immunofluorescence

hAEC cultures were fixed and stained for immunofluorescence as previously described (19). Well-differentiated cultures were stained using the following primary and secondary antibodies (tables 1 and 2).

**Table 1.** Overview of primary antibodies used in the current study

| 1° Antibody | Target | Dilution | Host | Clone | Supplier |
| --- | --- | --- | --- | --- | --- |
| Anti-β-Tubulin | Cilia | 1:200 | Mouse | ONS 1A6 | Abcam |
| Anti-ZO-1 | Tight junctions | 1:200 | Goat | Ab99462 |  |
| Anti-eGFP | eGFP | 1:200 | Mouse | Ab1281 |  |
| Anti-mCherry | mCherry | 1:200 | Chicken | Ab205402 |  |
| Anti-ZO-1 | Tight Junctions | 1:200 | Rabbit | 61-7300 | Thermofisher |

**Table 2.** Overview of secondary antibodies used in the current study

| 2° Antibody | Target | Dilution | Host | Supplier |
| --- | --- | --- | --- | --- |
| Alexa Fluor® 488 | Anti-mouse | 1:400 | Donkey | Jackson Immunoresearch |
| Cy3 | Anti-goat | 1:400 |  |  |
| Alexa Fluor® 647 | Anti-goat | 1:400 |  |  |
| Alexa Fluor® 594 | Anti-chicken | 1:400 |  |  |
| Alexa Fluor® 647 | Anti-rabbit | 1:400 |  |  |

All samples were counterstained with DAPI (4’,6-diamidino-2-phenylindole; Invitrogen, Fisher Scientific AG, Reinach, Switzerland) to visualize nuclei. Images were acquired on a Nikon confocal microscope A1 (Nikon GmbH, Egg, Switzerland) combined with an ECLIPSE Ti inverted microscope using a Plan Apo 60x/1.40 oil objective. Image capture, analysis and processing were performed using the Nikon (NIS-Elements AR 3.30.02) and Imaris 8.0.2 (Bitplane AG, Zurich, Switzerland) software packages.

### 2.10 Cell viability

Cell viability was assessed with Alamar Blue (Thermo Fisher Scientific). Alamar Blue was added to growth medium at 10% (v/v) end concentration and incubated at 37°C and 5% CO_2_ for at least 4 hours, depending on cell density. After incubation, fluorescence was measured in a luminometer at 595 nm.

### 2.11 Virus infection

Transgenic and naïve hAEC cultures were inoculated apically with 10.000 plaque forming units (PFU) of either human coronavirus 229E (HCoV-229E GFP [27]) or human respiratory syncytial virus (RSV-B GFP, [28]; kindly provided by Prof Dr. Paul Duprex, Boston University, School of Medicine) and incubated for 2 hours at 33°C in a humidified incubator with 5% CO_2_. Subsequently, inoculum was removed, and the apical surface washed three times with HBSS, after which the cells were incubated for 72 hours, with progeny virus collection every 24 hours by incubating 100 μL of HBSS on the apical surface for 10 minutes prior to collection. In parallel, basolateral medium supplemented with 0 or 2 mM IPTG was replaced every 24 hours. Progeny virus collections were stored 1:1 in virus transport medium (VTM) for later quantification [19].

### 2.12 eGFP knockdown

For eGFP knockdown in well-differentiated dually transduced hAEC cultures, cells were treated with 0 or 2 mM IPTG in basolateral medium over a period of 6 days, with media change every 24 hours. eGFP expression was analyzed on transcriptional level with qPCR, and on protein level via immunofluorescence and flow cytometry at the end of the treatment period.

### 2.13 Quantitative Real-time PCR (qRT-PCR)

Total cellular RNA from well-differentiated hAECs was extracted with the Nucleospin™ RNA extraction kit (Macherey-Nagel, Oensingen, Switzerland) according to the manufacturer’s guidelines. Reverse transcription was performed with GoScript™ reverse transcriptase mix random hexamers according to the manufacturer’s protocol (A2800; Promega AG, Dübendorf, Switzerland) using 200 ng of total RNA. Two microliters of tenfold diluted cDNA was amplified using Fast SYBR™ Green Master Mix (Thermo Fisher Scientific) according to the manufacturer’s protocol using primers targeting 18S and MxA as described previously [12]. Relative gene expression was calculated using the 2-ΔΔCt method [29] and is shown as fold induction over untreated controls.

For quantification of progeny virus in apical washes from HCoV-229E and RSV infected hAECs, viral RNA was extracted using the NucleoMag VET (Macherey-Nagel) according to the manufacturer’s instructions on a Kingfisher Flex Purification system (Thermo Fisher Scientific). Two microliters of extracted RNA was amplified using TaqMan™ Fast Virus 1-Step Master Mix (Thermo Fisher Scientific) according to the manufacturer’s protocol and primers specific for HCoV-229E [30] or RSV-B [31], or eGFP forward primer 5’-GGG CAC AAG CTG GAG TAC AAC −3’ and reverse primer 5’-CAC CTT GAT GCC GTT CTT CTG −3’. Measurements and analysis were performed using an ABI7500 instrument and software package (Applied Biosystems, Fisher Scientific AG, Reinach, Switzerland).

### 2.14 Data presentation

Data was plotted using GraphPad Prism 7 and figures were assembled in Adobe Illustrator CC 2018 software package.

## 3. Results

### 3.1 Transduction of undifferentiated primary human bronchial cells

In an initial effort to establish transgenic hAEC cultures, we attempted to transduce well-differentiated cultures directly with vesicular stomatitis virus glycoprotein (VSV-g) pseudotyped lentiviral particles harboring a pLKO_GFP transfer vector. However, intact cultures are refractory to both apical and basolateral transduction while damaged cultures can be transduced from the apical side when the basolateral side of the epithelium is exposed. In those cultures, GFP expression is only observed around the edges of damaged epithelium (data not shown). This is consistent with the observation that receptors for VSV-g are predominantly located on the basolateral side of polarized epithelia [32]. Since well-differentiated hAEC cultures cannot be efficiently transduced, we assessed the optimal duration and method of transduction of human bronchial cells in their undifferentiated state with VSV-g pseudotyped lentiviral particles harboring the GFP-modified pLKO backbone containing a non-mammalian shRNA control that should not target any known mammalian gene (pLKO_GFP_Scr), but will engage with the RNA-Induced Silencing Complex (RISC). We observed that GFP expression in undifferentiated bronchial cells generally varied between experiments. However, suspension transduction consistently resulted in the highest percentage of GFP-positive cells. Further modifications of the protocol such as the addition of polybrene, separately or in combination with spinoculation, did not seem to increase the efficacy of transduction (data not shown), leading to the final optimized protocol that utilizes suspension transduction for 4 hours, followed by additional 24-hour incubation of lentivirus containing supernatant during cell attachment in monolayer. Specifically, cells and lentiviral supernatant were incubated in suspension at 37°C and 5% CO_2_ with gentle shaking every hour for 4 hours, after which the mixture was seeded in T25 culture flasks in 4 mL total Bronchial Epithelial Growth Medium (BEGM). Medium was changed after 24 hours, bringing the total lentiviral incubation time to 28 hours. Using this protocol, we were able to transduce primary bronchial cells with an efficacy of 30-70%, depending on the donor and MOI used.

Since the distinctive cellular composition of well-differentiated hAEC cultures is essential to all virus – host interaction studies, the morphology of transgenic and naïve cultures must be interchangeable to preserve the functionality of the culture system. Using the established transduction protocol, we first evaluated whether hAECs differentiated normally after lentiviral transduction using flow cytometry. The cellular composition of heterogeneously transgenic hAECs that have the pLKO_GFP_Scr cassette integrated in their genome does not differ from naïve cultures, as measured by the division into ciliated (tubulin), goblet (Mucin1) and basal cells (NGFR) (Figure 1A). More importantly, GFP expression can be observed in all cellular subgroups within the differentiated cultures (Figure 1B). This data indicates that lentiviral transduction does not interfere with the differentiation potential of primary human bronchial cells and, importantly, that transgene expression (GFP) can still be observed 6 weeks post-differentiation using a fluorescent microscope (Figure 1C).

**Figure 1.**
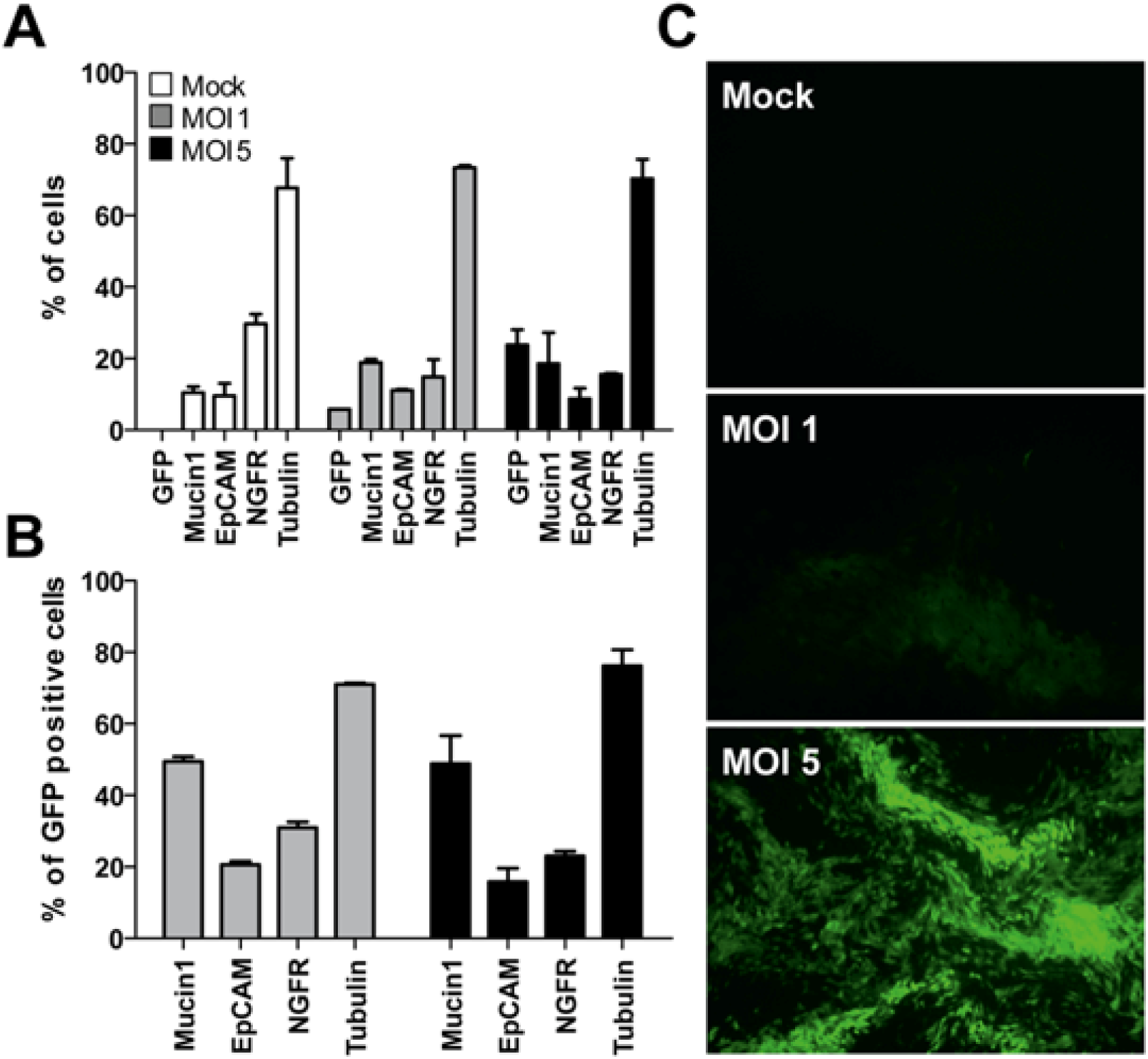
Transduction of primary airway epithelial cells does not alter cellular composition and differentiation. (**a**) Comparison of the cellular composition between naïve and heterogeneously transgenic hAECs transfuced with two different MOIs of lentivirus 6 weeks post-differentiation by FACS using cell-type specific markers for ciliated (tubulin), goblet (Mucin1), basal (NGFR) and transgene positive cells (GFP). (**b**) Transgene expression among different cell types. (**c**) A representative fluorescent microscopy image of GFP expression levels among naïve and transgenic hAECs 6 weeks post-differentiation.

Thus, we have shown that the cellular composition of transgenic cultures does not differ from naïve controls and GFP transgene expression can be observed in all cellular subgroups after 6 weeks of differentiation. Therefore, we conclude that we are able to successfully transduce progenitor basal cells in suspension prior to the generation of well-differentiated hAEC cultures. This suggests that FACS sorting of transgene positive cells would not deplete any cell population from the differentiated cultures and allow for the establishment of transgenic cultures with proper cellular composition and morphology.

### 3.2 Treatment of primary cells with Y-27632 prolongs basal cell phenotype

Suspension transduction of primary human tracheobronchial cells requires at least two additional passages prior to the establishment of homogenously transgenic hAEC cultures compared to naïve cells. After transduced cells have been expanded in monolayer, they must be sorted by flow cytometry and re-expanded. Only after that, can the cells be seeded on porous inserts and liquid-liquid interface (LLI) established. Once confluent, the apical medium is aspirated, establishing ALI. By incorporating the necessary transduction, expansion and sorting steps, transgenic tracheobronchial cells cannot be seeded in LLI before monolayer passage three. However, since differentiated hAEC cultures cannot be reliably established after monolayer passage two due to cellular senescence and loss of differentiation potential [16] we had to devise a method that would prolong the lifespan of primary human cells in monolayer culture to allow for lentiviral transduction, expansion and eventual cell sorting.

Treatment of both bronchial and cervical epithelial cells with a commercially available Rho-associated protein kinase (ROCK) inhibitor, Y-27632, has been shown to induce a basal epithelial cell phenotype, thereby preserving the differentiation capabilities of human epithelial cells in vitro [18]. We therefore assessed the differentiation potential of Y-27632 treated primary human bronchial cells in our ALI conditions. Indeed, when these cells were treated with 10 μM of Y-27632 they showed increased expression of alpha6-integrin and neural growth factor receptor (NGFR), common basal cell markers, compared to non-treated cells in later passage (Figure 2A). The up-regulation of these basal cell markers indicates that the epithelial basal cell phenotype is induced in treated cells, prolonging their life span and differentiation capabilities in vitro. Furthermore, hAEC cultures established with treated cells showed the same cellular composition as non-treated cells up to passage (p) 4, as measured by the expression of tubulin (ciliated cells), Mucin 1 (goblet cells), and NGFR (basal cells) (Figure 2B). Thus, the removal of the inhibitor from culture medium upon LLI culture seems to be sufficient for the cells to proceed through differentiation normally post-treatment. To confirm proper epithelial structure and morphology, hAEC cultures from both treated and non-treated cells were stained for nuclei (DAPI), cilia (β-tubulin) and tight junctions (ZO-1). In low passage (p2) the two groups share the same structure and morphology (data not shown). However, in p4, only Y-27632 treated cells, independent of donor, were able to maintain epithelial integrity post-differentiation. Non-treated cells in p4 seem to differentiate normally early on, evidenced by the presence of ciliated cells, but eventually the epithelial layer dissociates and detaches (Figure 2C).

**Figure 2.**
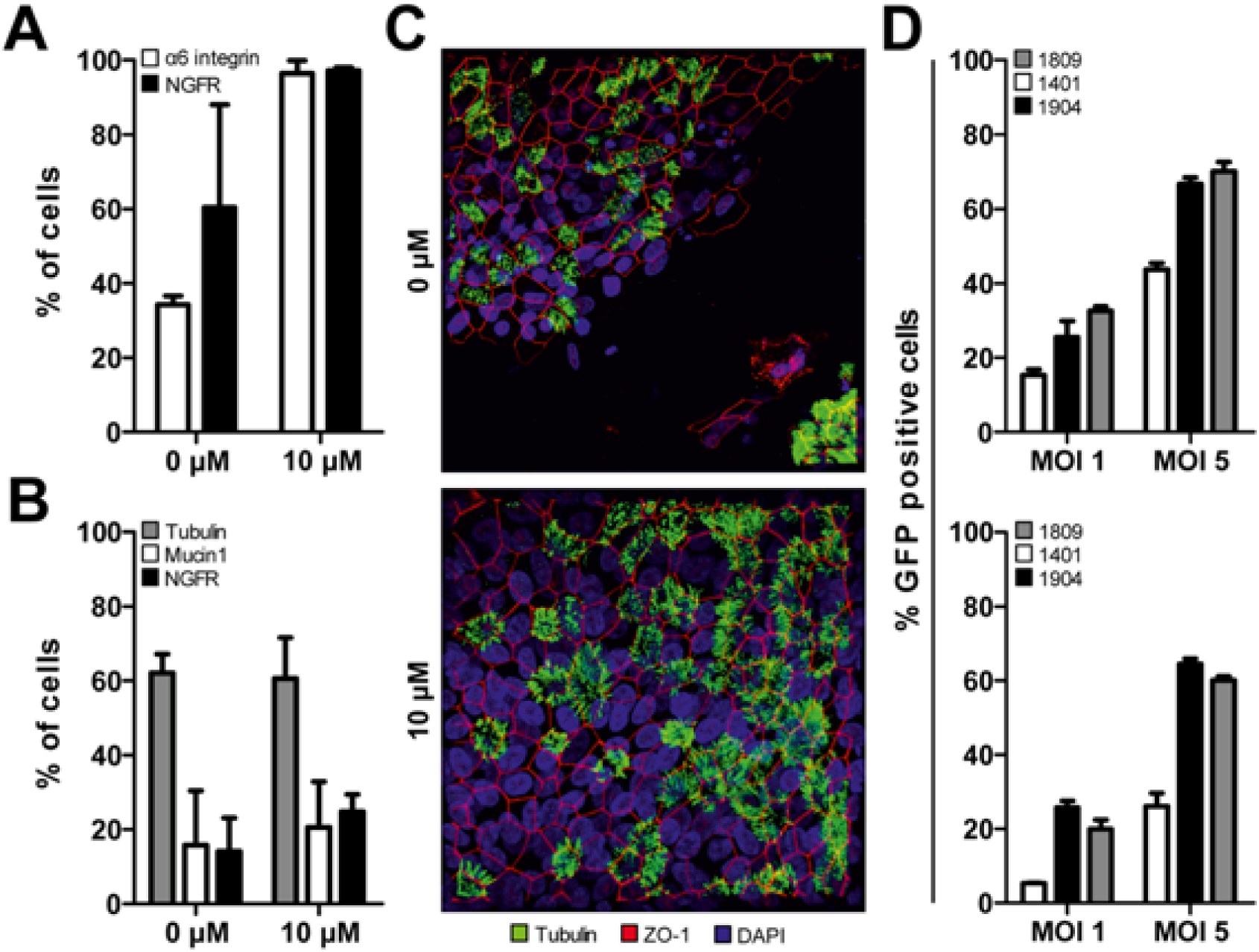
Treatment of primary human tracheobronchial epithelial cells with a Rho-associated protein kinase inhibitor (Y-27632) induces a basal cell phenotype and prolongs culture time. (**a**) Late passage (p4) monolayer hAEC cultures treated with 10 µM Y-27632 retain a basal cell phenotype. Data is shown as two technical replicates from two individual biological donors. (**b**) Naïve and Y-27632 treated well-differentiated hAEC cultures exhibit the same cellular composition, indicating that treatment of cells with Y-27632 does not affect cellular differentiation capabilities. Data is shown as mean of two technical replicates from two individual biological donors. (**c**) A representative maximum intensity Z-stack projection from late passage (p4) hAEC cultures treated with 0 or 10 µM Y-27632. Demonstrating that hAEC cultures treated with 10 µM Y-27632 are able to maintain epithelial integrity post-differentiation. Original magnification 63x. (**d**) Monolayer hAEC cultures treated with 10 µM of Y-27632 (bottom) can be transduced with lentiviral vectors to similar levels as non-treated hAEC culture (top). Data is shown as mean of two technical replicates for each individual biological donor.

Next, we assessed whether Y-27632 treated cells could be transduced to similar levels as naïve cells. Suspension transduction of cells isolated from different donors with an MOI of 1 or 5 results in varying levels of transduction efficacy (15-35% and 40-70%, respectively) evaluated by the percentage of GFP positive cells (Figure 2D, top). Treatment of primary tracheobronchial cells with Y-27632 results in slightly lower transduction efficacies under the same conditions (5-30% and 30-60%, figure 2D, bottom). Interestingly, even when a basal phenotype has been induced in all three donors the donor variability is still present and slightly exaggerated compared to naïve cultures (Figure 2D). These results indicate that the modification of our protocol, namely, treatment of primary human bronchial cells with Y-27632, induces a prolonged basal cell phenotype in our primary human bronchial cells. This will accommodate the additional passages required for cellular expansion post-lentiviral transduction and cell sorting to establish homogenously transgenic hAEC cultures.

### 3.3 Establishment of homogenously transgenic hAEC cultures

Thus far, we have shown that the establishment of homogenously transgenic hAEC cultures is feasible, provided changes are made to the standard cell culture protocol. However, since the differentiation of hAEC cultures is a delicate process, constitutive knockdown of certain genes could possibly interfere with differentiation resulting in unusable hAEC cultures post-transduction. For example, the knockdown of p63, a basal cell transcription factor, results in the loss of epithelial integrity in hAEC cultures established from a cell line while p63 knockdown in primary cells results in accelerated cellular senescence and death before ALI can be established [33]. To circumvent these limitations, we aimed to establish an inducible lentiviral system for gene knockdown in hAEC cultures. To this end, we modified the inducible shRNA expression lentiviral vector pLKO_Puro_IPTG_3xLacO to constitutively express the mCherry fluorescence gene instead of a puromycin selection marker, hereafter referred to as pLKO_mCherry_3xLacO. This allows for cell sorting and microscopic evaluation of cells transduced with this shRNA cassette. Treatment of transduced cells with Isopropyl β-D-1-thiogalactopyranoside (IPTG), a lactose mimic, induces the expression of the shRNA controlled by the Lactose operon (LacO). For initial evaluation and optimization of the generation of homogenously transgenic hAEC cultures, we incorporated the non-mammalian shRNA (shSCR) in our inducible lentiviral vector (pLKO_mCherry_3xLacO).

To establish homogeneously transgenic cultures, human tracheobronchial cells were transduced using the previously established suspension transduction protocol. After initial expansion, undifferentiated primary airway epithelial cells were sorted for mCherry-positive cells by FACS. The mCherry-positive cells were then further expanded prior to the establishment of homogenously transgenic hAEC cultures. Comparison of naïve and transgenic hAEC cultures revealed the same morphology, a pseudostratified layer with distinct ciliated cells (β-tubulin, green) and tight junctions (ZO-1, cyan). Furthermore, nuclear expression of mCherry was observed in all nuclei in the epithelial layer (red) (Figure 3). This confirms our previous hypothesis that transduction and FACS sorting of undifferentiated primary bronchial cells does not deplete any cell population from the resulting differentiated cultures and allows for the establishment of homogenously transgenic hAEC cultures with correct anatomical morphology and uniform transgene distribution.

**Figure 3.**
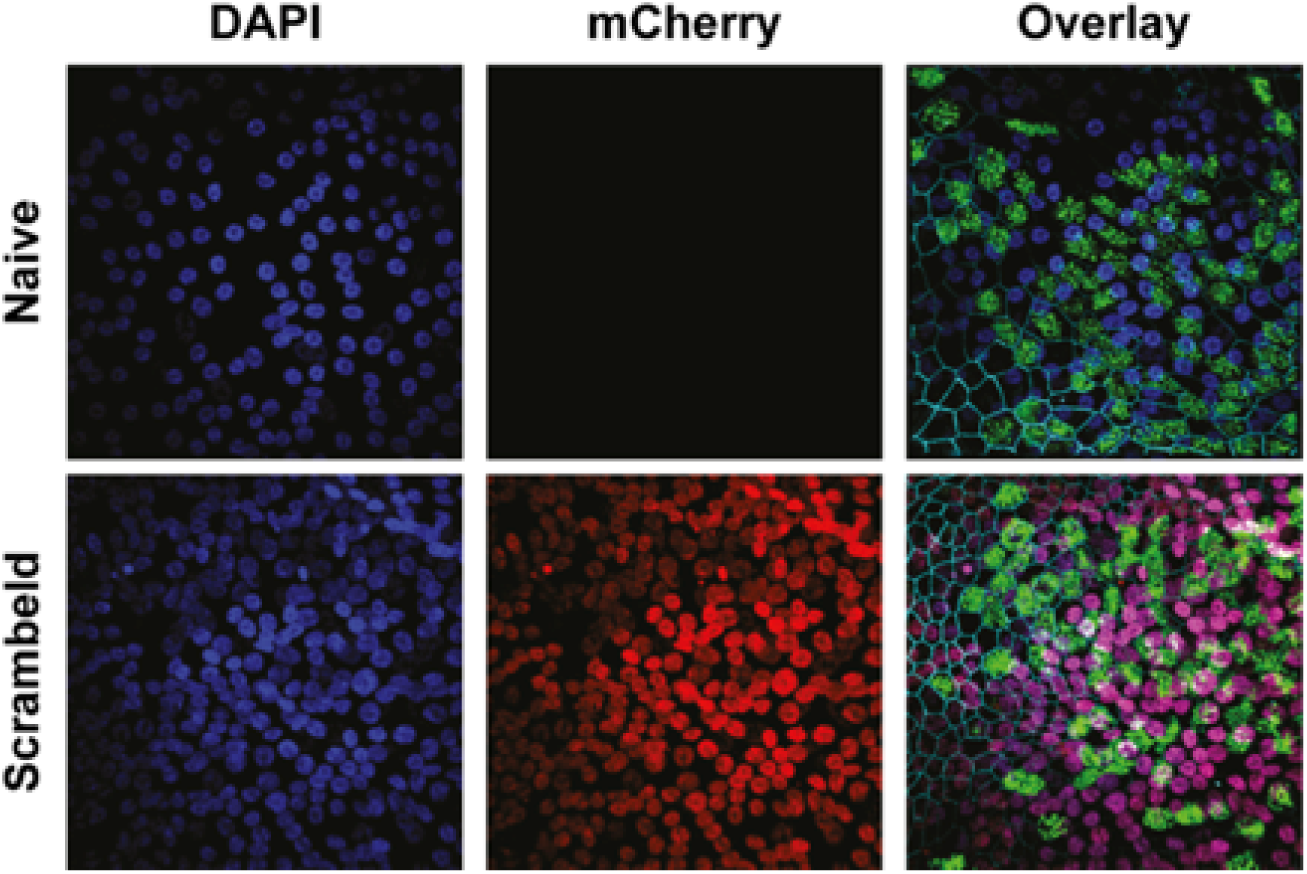
Comparison of the morphology of naïve and homogeneously transgenic hAEC cultures. Naïve and transgenic hAEC cultures were formalin-fixed 6 weeks post-differentiation and immunostained with antibodies to visualize the cilia (β-tubulin IV, green), tight junction borders (ZO-1, cyan), mCherry (red) and counterstained with DAPI to visualize the nuclei (Blue). A representative maximum intensity Z-stack projection of two individual biological donor is displayed and reveals correct anatomical morphology and uniform transgene distribution homogeneously transgenic hAEC cultures. Original magnification 63x.

### 3.4 Successful induction of IPTG-controlled shRNA expression

After having successfully demonstrated that homogenously transgenic hAEC cultures differentiate normally after lentiviral transduction, we generated two additional pLKO_mCherry_3xLacO vectors containing shRNAs targeting the CDS of the GFP mRNA transcript, shGFP1 and shGFP2, respectively. We first evaluated the effectiveness of these constructs in conventional Huh7 cells, as the establishment of homogenous transgenic hAEC cultures requires 6-8 weeks. After simultaneous transduction with two different lentiviral vectors, pLKO_GFP and pLKO_mCherry_3xLacO, containing shGFP1, shGFP2 or shSCR, the cells were sorted for double positive GFP/mCherry signal. When we treated double positive cells with 1 mM IPTG for 72 hours, a reduction in the number of GFP-positive cells could be observed in cells transduced with shGFP, but not in those with shSCR (figure 4A, top). The reduction of the total number of GFP-positive cells does not exceed 30% for either GFP specific shRNA. However, when comparing the median fluorescence intensity (MFI), there is 60-70% reduction of GFP fluorescence in shGFP cells while shSCR cells have an MFI similar to non-treated controls (figure 4A, bottom). In accordance, fluorescence microscopy also shows a prominent reduction of fluorescence intensity in treated cells compared to control (Figure 4B). The knockdown of GFP in Huh7 cells upon induction with IPTG demonstrates the functionality of different shRNA constructs in our modified inducible pLKO_mCherry_3xLacO lentiviral vector.

**Figure 4.**
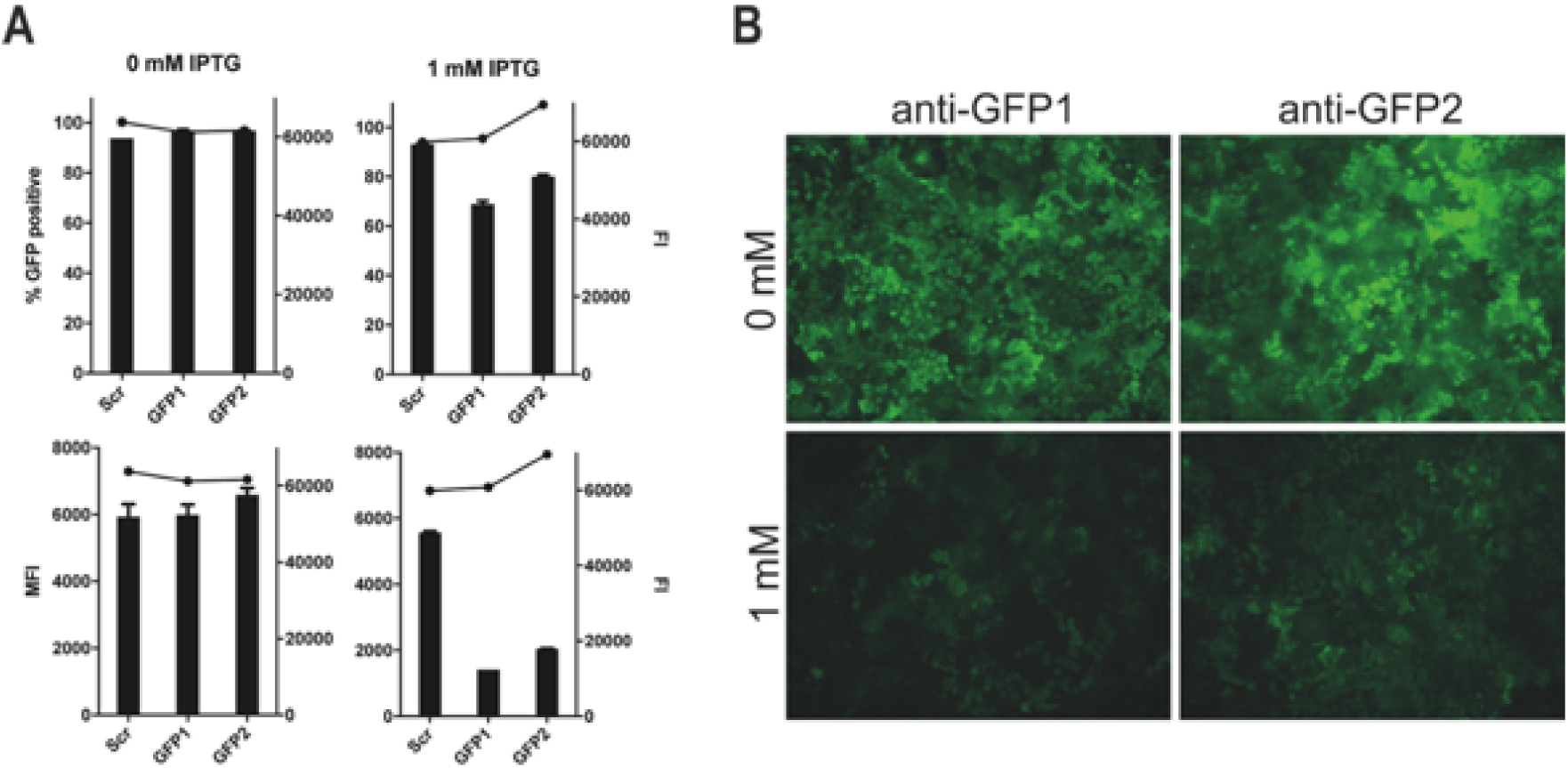
Down-modulation of GFP with an IPTG-inducible shRNA. (**a**) Huh-7 cells were treated with 0 or 1 mM of IPTG for 72 hours after which the percentage (top left y-axis) and mean fluorescence intensity (MFI; bottom left y-axis) GFP expression was assessed by FACS, in parallel the cell viability was assessed with Alamar Blue (Fluorescence Intensity (FI); right y-axis). The results are shown as means and SD from duplicates from one independent experiment. (**b**) A representative microscopic image displaying the GFP expression of Huh-7 cells harboring either the shGFP1 and shGFP2 construct after 72 hours of treatment with 0 or 2 mM IPTG. Original magnification 10x.

**Figure 5.**
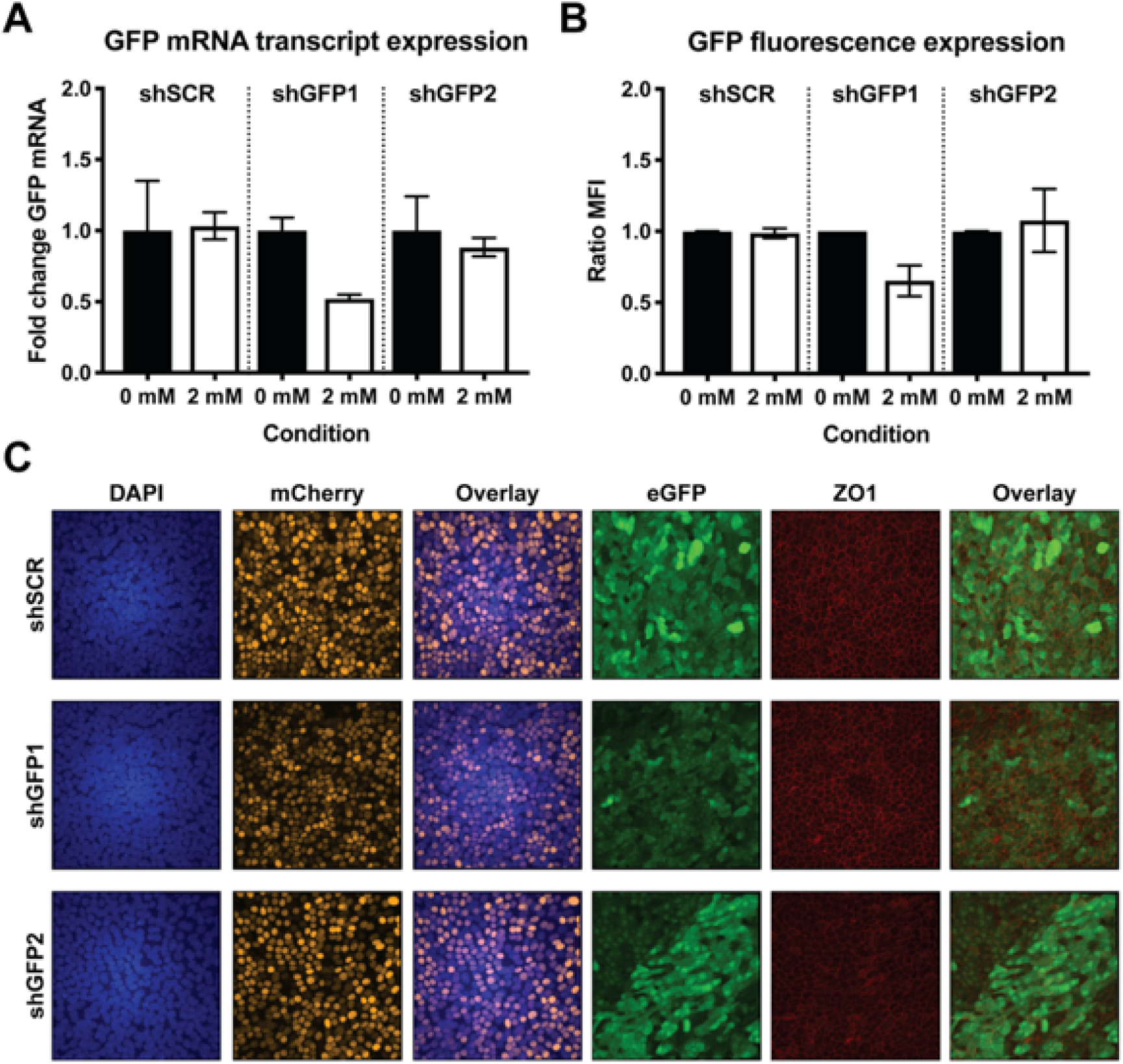
Down-modulation of GFP in well-differentiated homogenous transgenic hAEC cultures. Six-week-old well-differentiated homogenous transgenic hAEC cultures were treated with 0 or 2 mM of IPTG for 6 consecutive days after which the GFP protein expression was analyzed by FACS (**a**) and the GFP mRNA expression by qPCR (**b**). The GFP protein expression data is shown as mean and SD from three independent biological donors, whereas the mRNA expression data is shown as mean and SD from two independent biological donors. (**c**) Moreover, the fluorescence intensity of GFP was assessed in formalin-fixed well-differentiated homogenous transgenic hAEC cultures after 6 days of 2 mM IPTG treatment. The cultures were immunostained with antibodies to visualize the mCherry (yellow), GFP (green) and tight junction borders (ZO-1, red) and counterstained with DAPI to visualize the nuclei (Blue). A representative maximum intensity Z-stack projection from two individual biological donors is displayed. Original magnification 63x.

To demonstrate that host gene expression can also be modulated post-differentiation via inducible shRNA-mediated knockdown we repeated these experiments in primary hAEC cultures using the same method of transduction and cell sorting as described above. After 6 weeks of differentiation, both transgenic and naïve cultures were maintained for 6 consecutive days in the presence or absence of 2 mM IPTG after which GFP expression was analyzed by flow cytometry. Upon induction, we observed approximately 40% reduction in MFI in hAEC cultures expressing shGFP1, whereas induction of shSCR and shGFP2 did not result in pronounced reduction of fluorescence (Figure 4A). This data correlates with the relative GFP mRNA expression level after IPTG-induction where a 50% reduction was observed for shGFP1 alone (Figure 4B). To further corroborate these findings, we used confocal microscopy to visualize GFP expression in transgenic hAEC cultures treated with 2 mM IPTG. Microscopic analysis showed homogenously transgenic hAEC cultures expressing both mCherry and GFP, as well as the characteristic pattern of the tight-junction marker ZO-1. Furthermore, we observed that GFP signal intensity is heterogeneously distributed in all cultures, but overall the GFP expression in shGFP1 cultures was distinctly weaker compared to shSCR and shGFP2 after induction (Figure 4C). These observations are in agreement with our previous results and provide additional validation of the functionality of our modified lentiviral vector as well as confirming that gene expression can be modulated in hAEC cultures post-differentiation via inducible shRNA-mediated knockdown. However, given the observed discrepancies in the effectiveness of shGFP2 between Huh7 cells and hAEC cultures, individual shRNA constructs intended for the modulation of gene expression in hAEC cultures require careful evaluation and testing.

### 3.5 Transgenesis of primary hAEC cultures does not affect host innate immune response or viral replication

In the airways, respiratory pathogens can be detected through the recognition of pathogen-associated molecular patterns (PAMPs) by pattern recognition receptors (PRRs) which are expressed in the hAEC cultures akin to human airway epithelium in vivo [34]. Polyinosinic:polycytidylic acid (poly(I:C)) is a synthetic analogue of double stranded RNA (dsRNA), a molecular pattern associated with viral infection that can be recognized by the PRRs TLR-3, RIG-I, and MDA5 [35,36]. The subsequent signaling cascade initiates the production of interferon (IFN) that leads to the induction of several hundred interferon stimulated genes (ISGs), such as the well characterized Myxovirus resistance protein 1 (MxA) [37]. In order to determine whether our genetic modification influences the host innate immune response in transgenic hAEC cultures, we treated both naïve and homogenously transgenic cultures (shGFP1, shGFP2 and shSCR) with exogenous poly(I:C), as a proxy for viral infection, after 6-day pre-treatment with 0 or 2 mM IPTG to induce both mRNA and protein knockdown of GFP. 24 hours post poly(I:C) treatment, we assessed the expression level of MxA mRNA transcripts, revealing similar gene expression levels in both naïve and transgenic cultures regardless of shRNA construct and IPTG treatment conditions (Figure 6A). This indicates that neither the genetic modification, nor the presence of IPTG influences the innate immune response in homogenously transgenic hAEC cultures.

**Figure 6.**
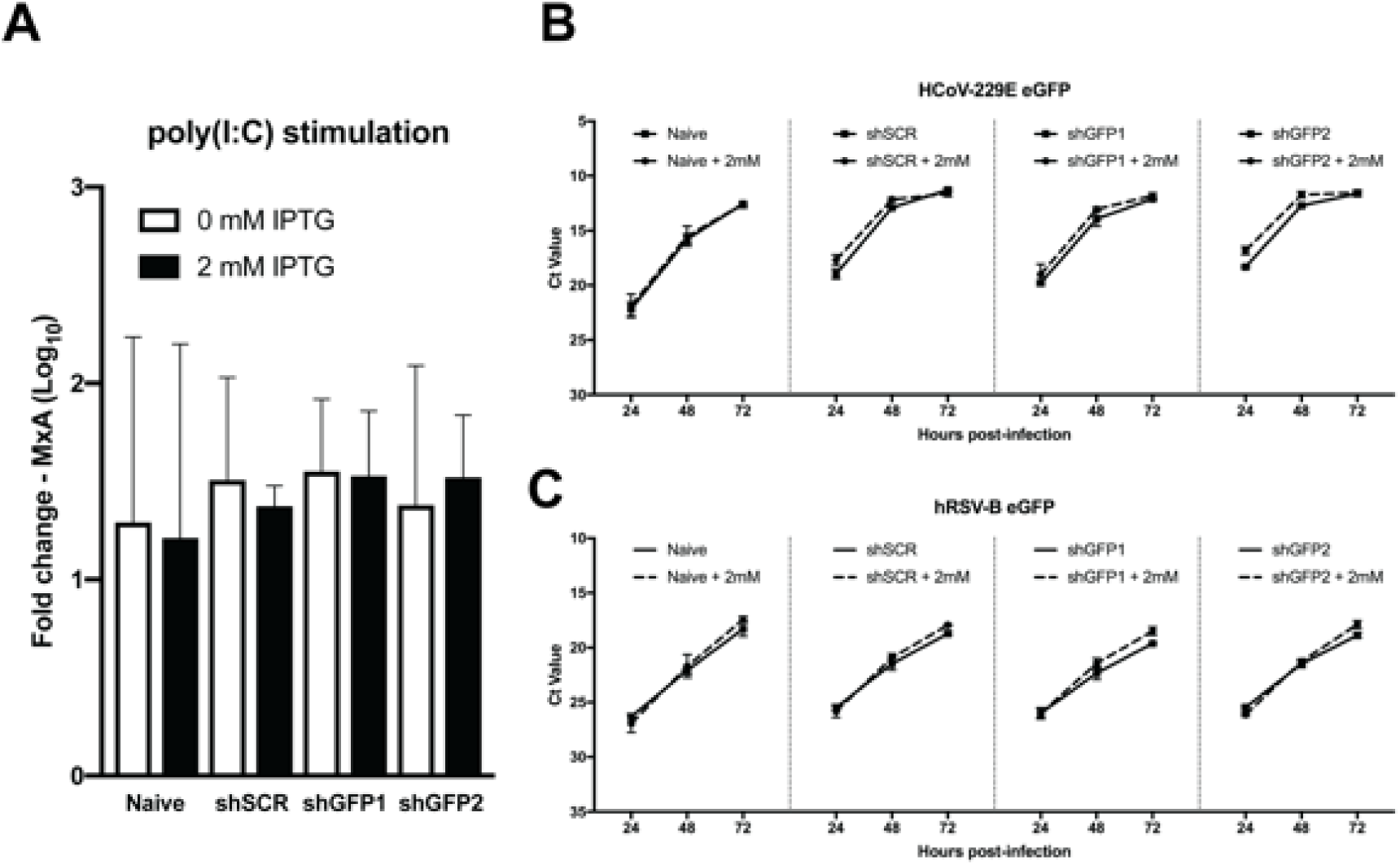
Transgenesis of primary hAEC cultures does not affect host innate immune response or viral replication. (**a**) Six-week-old well-differentiated naïve and homogenous transgenic hAEC cultures were for 6 consecutive days treated with 0 or 2 mM of IPTG. Afterwards hAEC cultures were stimulated for an additional 24 hours with exogenous poly(I:C) after which the fold change in MxA mRNA expression was assessed via qPCR. Data is shown as mean and SD from duplicates from two individual biological donor. (**b**) Alternatively, 0 and 2 mM IPTG-treated naïve and homogenous transgenic hAEC cultures were inoculated with 10.000 PFU of HCoV-229E-GFP or hRSV-B-GFP and incubated at 33°C. The monitored viral RNA yield is given as Cycle-threshold (Ct) value of isolated RNA (y-axis) at indicated hours post-inoculation (x-axis) for 33°C. Data is shown as mean and SD from triplicates from two individual biological donors.

In light of these results, we evaluated whether viral replication could be influenced by shRNA-mediated knockdown during hAEC infection with GFP expressing human coronavirus 229E (HCoV-229E-GFP). As stated previously, both naïve and homogenously transgenic hAEC cultures were pre-treated with 0 or 2 mM IPTG for 6 days prior to inoculation with 10.000 PFU of HCoV-229E-GFP. Quantification of viral RNA in apical wash revealed similar replication kinetics for all conditions, indicating that general transgenesis of hAEC cultures does not influence the replication of HCoV-229E-GFP (Figure 6B). However, unfortunately we did not observe any specific reduction in replication in those cultures were GFP-directed shRNA had been induced by IPTG (Figure 6B). Since we have demonstrated that our modified pLKO_mCherry_3xLacO lentiviral vector is functional and able to reduce the expression of cellular GFP upon IPTG induction of GFP-directed shRNAs, we speculated the failure of GFP knockdown to be cell type specific since HCoV-229E predominantly infects non-ciliated cells [13]. Therefore, we repeated the experiment with a GFP-expressing human respiratory syncytial virus (hRSV-GFP) which exhibits a ciliary cell tropism [28]. The obtained results were in accordance with our previous observations, no difference in viral kinetics between naïve and homogenously transgenic hAEC cultures was observed after infection with hRSV-GFP. Moreover, as with HCoV-229E-GFP, we did not observe any reduction of viral replication kinetics in induced cultures expressing GFP-directed shRNAs (Figure 6C). This suggests that in the context of our experiments, neither HCoV-229E-GFP nor hRSV-GFP are susceptible to shRNA-mediated knockdown. However, it also indicates that our results are not cell type dependent, further asserting the functionality of our transgenic system as a whole. Based on these observations, we rather hypothesize that the lack of knockdown is due to an insufficient amount of available shRNA within infected cells to negatively influence the exponential increase of viral GFP mRNA expression while the levels of cellular GFP mRNA are more suitable for shRNA-mediated knockdown.

## 4. Discussion

In this study, we developed a robust and reproducible protocol to generate transgenic hAEC cultures. We demonstrate that the transduction of primary human tracheobronchial airway epithelial cells with lentiviral vectors in their undifferentiated state does not interfere with the cellular composition of the resulting well-differentiated airway epithelial cell cultures. Furthermore, we have shown that incorporation of the Rho-kinase associated inhibitor Y-27632 during the expansion phase induces a basal cell phenotype and increases the longevity of primary human bronchial cells. These results are in direct agreement with the initial publications that first described the induction and prolongation of a basal cell phenotype in primary cells without gross influence on cell differentiation capacity [17,18]. By incorporating Y-27632 during the cell propagation phase, we increased the number of attainable passages prior to the generation of well-differentiated hAEC cultures. In parallel, by optimizing our lentiviral transduction procedure, we could reproducibly achieve a transduction efficacy between 30 - 70% in undifferentiated bronchial cells, depending on the amount of lentivirus and donor used. This allowed us to incorporate a constitutive fluorescent reporter gene into the host cell genome that facilitates fluorescence-activated cell sorting (FACS) and the subsequent generation of homogenously transgenic hAEC cultures that differentiate normally. This fundamental change in our primary airway epithelial cell culture protocol has been pivotal to successful cellular expansion post-lentiviral transduction and, subsequently, the establishment of transgenic hAEC cultures suitable for virus – host interaction studies. During the development and validation of our protocol other research groups have reported alternatives or improvements to the use of the Rho-kinase associated inhibitor to further extend the life-span of primary human bronchial cells [38–41]. We anticipate that the incorporation of these alternative inhibitors and methods will further improve the currently established protocol to generate homogenously transgenic hAEC cultures, and might even facilitate the adaptation of this protocol to primary airway epithelial cells from other species to further elucidate molecular virus – host interactions without using animal models.

In the current study, we have demonstrated that we can modulate gene expression in transgenic hAEC cultures post-differentiation via inducible shRNA-mediated knockdown. However, gene expression modulation using shRNA requires careful evaluation of the effectiveness of individual shRNA constructs, as only one out of two shRNAs against GFP was effective in our homogenously transgenic hAEC cultures while both were effective in a cell line model. Additionally, we observed that GFP brightness was reduced by approximately 40% upon induction, and in the context of modulating host gene expression to study virus – host interactions this might not be sufficient to observe a phenotype. Nonetheless, since we demonstrate that an inducible lentiviral system is operational post-differentiation, alternative methods of host gene modifications can now be explored. For instance, microRNA (miRNA) mimics in conjunction with a fluorescent reporter gene under the control of an inducible Polymerase II promoter [42] would allow monitoring and quantification of the expression level of the miRNA mimic, providing a better-controlled system. Alternatively, employing inducible CRISPR-mediated gene editing would allow the modulation of host gene expression by either repression (CRISPRi) or activation (CRISPRa) [43]. All these potential methods require lentiviral-mediated transduction, which can be achieved with the currently established protocol.

Unfortunately, in our experiments, we did not observe any inhibition of viral replication through shRNA-mediated knockdown targeting the GFP gene in the HCoV-229E and hRSV reporter viruses. Since we observed the same phenotype with two viruses with differential cell tropism, non-ciliated and ciliated cells respectively, there seems to be no cell-type bias for shRNA-mediated knockdown in homogenously transgenic hAEC cultures. This is further corroborated by the uniform reduction of GFP expression by shGFP1 in our double transgenic hAECs, in which the MFI of GFP fluorescence is reduced by approximately 40%. Due to low inoculation dose (MOI 0.1), we hypothesize that the amount of shRNA targeting the viral GFP mRNA transcript simply is not sufficient to achieve a pronounced reduction of the logarithmic increase of viral replication. Adaptation of the inoculation dosage or even the Polymerase III promoter in the lentiviral vector to achieve a higher cellular concentration of shRNA might reveal a different phenotype. It has been reported that the viral nucleoprotein of HCoV-229E counteracts RNA silencing [44], which might also be the case for hRSV, rendering our shRNA ineffective against these viruses. Further characterization and validation of this phenomenon is fascinating but beyond the scope of this study.

Taken together, these results demonstrate that we have successfully established a robust and reproducible protocol to make hAEC cultures amenable to genetic modification using lentiviral vectors. This will greatly facilitate detailed studies on molecular virus – host interactions in primary human airway epithelium.

## Author Contributions

Conceptualization, Hulda Jonsdottir, Volker Thiel and Ronald Dijkman; Data curation, Hulda Jonsdottir and Ronald Dijkman; Formal analysis, Hulda Jonsdottir and Ronald Dijkman; Funding acquisition, Volker Thiel and Ronald Dijkman; Investigation, Hulda Jonsdottir and Ronald Dijkman; Methodology, Hulda Jonsdottir, Sabrina Marti and Ronald Dijkman; Resources, Dirk Geerts and Regulo Rodriguez; Supervision, Volker Thiel and Ronald Dijkman; Visualization, Hulda Jonsdottir and Ronald Dijkman; Writing – original draft, Hulda Jonsdottir; Writing – review & editing, Hulda Jonsdottir and Ronald Dijkman.

## Acknowledgments

We acknowledge Prof. Dr. B. Berkhoud and Dr. J. Eekels for providing lentiviral plasmids for lentiviral particle production, Prof. Dr. P. Duprex for contributing GFP expressing Respiratory Syncytial Virus, and Dominik Florek for his efforts during the initial stages of the project.

